# How to deal with toxic amino acids: the bipartite AzlCD complex exports histidine in *Bacillus subtilis*

**DOI:** 10.1101/2022.07.07.499250

**Authors:** Janek Meißner, Thorben Schramm, Ben Hoßbach, Katharina Stark, Hannes Link, Jörg Stülke

## Abstract

In the Gram-positive model bacterium *Bacillus subtilis*, the presence of the amino acid glutamate triggers potassium uptake due to the glutamate-mediated activation of the potassium channel KtrCD. As a result, the intracellular accumulation of glutamate is toxic in strains lacking the second messenger cyclic di-AMP since these cells are unable to limit potassium uptake. We observed that the presence of histidine, which is degraded to glutamate, is also toxic for a *B. subtilis* strain that lacks all three c-di-AMP synthesizing enzymes. However, suppressor mutants emerged, and whole genome sequencing revealed mutations in the *azlB* gene encoding the repressor of the *azl* operon. This operon encodes an exporter and an importer for branched-chain amino acids. The suppressor mutations result in overexpression of the *azl* operon. Deletion of the *azlCD* genes encoding the branched-chain amino acid exporter restored the toxicity of histidine indicating that this exporter is required for histidine export and resistance to otherwise toxic levels of the amino acid. The higher abundance of the amino acid exporter AzlCD increased the extracellular concentration of histidine, thus confirming the new function of AzlCD as a histidine exporter. Unexpectedly, AzlB-mediated repression of the operon remains active even in the presence of amino acids suggesting that expression of the *azl* operon requires mutational inactivation of AzlB.

**IMPORTANCE:** Amino acids are building blocks for protein biosynthesis in each living cell. However, due to their reactivity as well as the similarity between several amino amino acids, they may also be involved in harmful reactions or in non-cognate interactions and thus be toxic. *Bacillus subtilis* can deal with otherwise toxic histidine by overexpressing a bipartite amino acid exporter AzlCD. Although encoded in an operon that also contains a gene for an amino acid importer, the corresponding genes are not expressed, irrespective of the availability or not of amino acids in the medium. This suggests that the *azl* operon is a last resort to deal with histidine stress that can be expressed due to mutational inactivation of the cognate repressor, AzlB.

## INTRODUCTION

Amino acids are the essential building blocks for protein biosynthesis and many other cellular components. Cells can acquire amino acids by uptake from the environment, by degradation of external peptides or proteins, or by *de novo* biosynthesis. Many bacteria such as the model organisms *Escherichia coli* and *Bacillus subtilis* can use all three possibilities for amino acid acquisition. Although amino acids are essential for growth, they can be toxic due to misloading of tRNAs resulting in misincorporation into proteins and from their high reactivity. Moreover, many amino acids are chemically very similar to each other, and one amino acid that is available in excess may competitively inhibit the biosynthetic pathway(s) of similar amino acids by binding to the corresponding enzymes (1).

We are interested in the identification of the components that allow life of a simple minimal cell and in the construction of such cells based on the model bacterium *B. subtilis* (2, 3). Such minimal organisms are not only important to get a comprehensive understanding of the requirements of cellular life but they are also important workhouses in biotechnological and biomedical applications. Indeed, minimal organisms have been proven to be superior to conventionally constructed strains in the production and secretion of difficult proteins and lantibiotics (4, 5, 6). For *B. subtilis*, the pathways for all amino acid biosyntheses have been completely elucidated. In contrast, the knowledge about amino acid transport is far from being complete as for several amino acids no transporter has been identified so far. This knowledge is important for the construction of genome-reduced strains that may be designed to grow in complex or minimal medium and thus require the complete set of uptake or biosynthetic enzymes, respectively. Moreover, some amino acids such as glutamate are toxic for *B. subtilis* even at the concentrations present in standard complex medium if the catabolizing enzymes, e.g. glutamate dehydrogenase, are absent (7, 8). Thus, a complete understanding of all components involved in cellular amino acid homeostasis is required for the succsessful generation of minimal organisms.

Amino acid toxicity is not only relevant for the design of minimal genomes but it is also an important tool for the identification of components involved in amino acid metabolism. While some amino acids such as serine or threonine are toxic already for wild type strains (1, 9, 10), others are well tolerated. In this case, the corresponding D-amino acids, amino acid analogs, or structually similar metabolites may act as anti-metabolites that inhibit normal cellular metabolism and thus growth of the bacteria. The application of toxic amino acids or of similar compounds to a bacterial growth medium will inhibit growth but will also result in the acquisition of suppressor mutations that allow the cells to resolve the issue of amino acid toxicity. Often, such mutations affect uptake systems and prevent the uptake of the toxic amino acid or its analogues. In this way, transporters for threonine, proline, alanine, serine, and glutamate as well as for the anti-metabolites 4-hydroxy-L-threonine and glyphosate have been identified in *B. subtilis* (9, 10, 11, 12, 13, 14, 15). A second way to achieve resistance against toxic amino acids is the activation of export mechanisms. This has been reported in the cases of 4-azaleucine and glutamate (14, 16). Third, suppressor mutations may facilitate detoxification of the toxic amino acid by degradation or modification to a non-toxic metabolite as observed for glutamate and serine (7, 10, 14, 17). Forth, the protein target of the toxic metabolite may be modified in a way that it becomes resistant (18). Finally, the bacteria can escape inactivation by increased expression of the target protein as has been reported for serine and the anti-metabolite glyphosate (10, 15).

Recently, it has been shown that the sensitivity of *B. subtilis* to glutamate is strongly enhanced if the bacteria are unable to produce the second messenger cyclic di-AMP (c-di-AMP) (14). This second messenger is essential for growth of *B. subtilis* on complex medium, and it is toxic upon intracellular accumulation (19). Both essentiality and toxicity are mainly a result of the central role of c-di-AMP in potassium homeostasis. The second messenger prevents the intracellular accumulation of potassium by inhibiting potassium import and by stimulating potassium export. Thus, the intracellular potassium concentration is kept within a narrow range (19, 20). The presence of high potassium concentrations in a strain lacking c-di-AMP results in the activation of potassium export by the acquisition of mutations in a sodium/H^+^ antiporter. These mutations change the specificity of the antiporter towards potassium (21). Even though none of the known targets of c-di-AMP is directly involved in glutamate homeostasis, glutamate is as toxic as potassium to the mutant lacking all diadenylate cyclases that would synthesize c-di-AMP. This can be explained by the fact that glutamate activates the low-affinity potassium channel KtrCD by strongly increasing the affinity of this channel. Thus, even the very low potassium concentration, which must be present even for the growth of this strain, become toxic due to the high affinity of KtrCD for potassium in the presence of glutamate (22). Accordingly, the *△dac* strain lacking c-di-AMP acquires mutations that reduce potassium uptake if propagated in the presence of glutamate. In addition, the bacteria usually acquire additional mutations that interfere with glutamate homeostasis by reducing uptake, facilitating export, or allowing degradation of the amino acid (14).

In this study, we were interested in the control of histidine homeostasis. Histidine biosynthesis from ribose 5-phosphate requires ten enzymes (see http://subtiwiki.uni-goettingen.de/v4/category?id=SW.2.3.1.14; 23). The degradation of histidine to ammonia, glutamate, and formamide involves a specific transporter, HutM, and four enzymes. The histidine transporter is induced in the presence of histidine, which is a typical feature for high-affinity transport systems (24, 25).Usually, high-affinity transporters are used for catabolic pathways to use an amino acid as carbon or nitrogen source. In contrast, constitutively expressed low-affinity transporters are used to import the amino acid from complex medium for protein biosynthesis. In many cases, both, low- and high-affinity amino acid transporters, are encoded in the genome of *B. subtilis*, and they are expressed depending on the pyhsiological conditions. However, no low-affinity histidine transporter has been identified so far. Histidine degradation yields intracellular glutamate, which is toxic for mutants lacking c-di-AMP (14) due to the activation of the potassium channel KtrCD (22). We thus expected that the strain lacking c-di-AMP has a similar sensitivity against histidine as it is against glutamate. We made use of this sensitivity to the degradation product glutamate to get further insights into the components that contribute to histidine homeostasis in *B. subtilis.* Our study revealed that mutational activation of an export system is the major mechanism to achieve resistance to histidine.

## RESULTS

### Histidine is toxic to a *B. subtilis* strain lacking c-di-AMP, and mutations in the *azlB* gene overcome the toxicity

Some amino acids such as serine and threonine are toxic for *B. subtilis.* In the case of glutamate, toxicity becomes visible in the absence of a degradation pathway or if the bacteria are unable to form the second messenger c-di-AMP (14). Here, we tested growth of a wild type strain (168) of *B. subtilis* and of an isogenic strain that had all three genes encoding diadenylate cyclases deleted (Δ*dac*, GP2222) (21) in the presence of histidine. As shown in Fig. 1, growth of the *B. subtilis* wild type strain 168 is not affected by histidine concentrations up to 10 mM. At a higher concentration of 20 mM, growth was inhibited. In contrast, histidine is highly toxic for *B. subtilis* GP2222 even at very low histidine concentrations (see Fig. 1). We observed that larger colonies rapidly appeared. It is likely that these larger colonies were formed bysuppressor mutants that were resistant to histidine in the medium. We hypothesized that mutations could affect uptake systems for histidine as already observed for glutamate, serine, or threonine (9, 10, 14). Indeed, we were able to indentify mutations in two isolates by whole genome sequencing. However, to our surprise, the mutations did not cover known or putative amino acid transporters of *B. subtilis* (23). In contrast, we observed mutations in the *azlB* gene, which encodes a Lrp-type transcription repressor that controls the expression of a branched chain amino acid exporter (AzlCD) and a branched chain amino acid importer (BrnQ) (9, 26). Strain GP3638 carried an amino acid substitution in AzlB (Asn24 Ser). In the second strain, GP3639, we found an eight basepair insertion (CATTAATG) after the 37^th^ basepair of the coding sequence that results in a frameshift and thus prevents the expression of a functional AzlB protein. As the *azlB* gene seemed to be a hotspot of mutations in histidine-resistant suppressor mutants, we determined the sequence of this gene in four additional mutants. In each case, mutations were present in the *azlB* gene, either amino acid substitutions, Asn24 Ser as in GP3638, Ile31 Met, or frameshift mutations. Since the frameshift mutations prevent the formation of functional AzlB proteins, it seemed likely that the amino acid substitutions also resulted in inactive proteins. Indeed both the N24S and the I31M mutations affect the DNA-binding helix-turn-helix motif of AzlB.

**Fig. 1.**
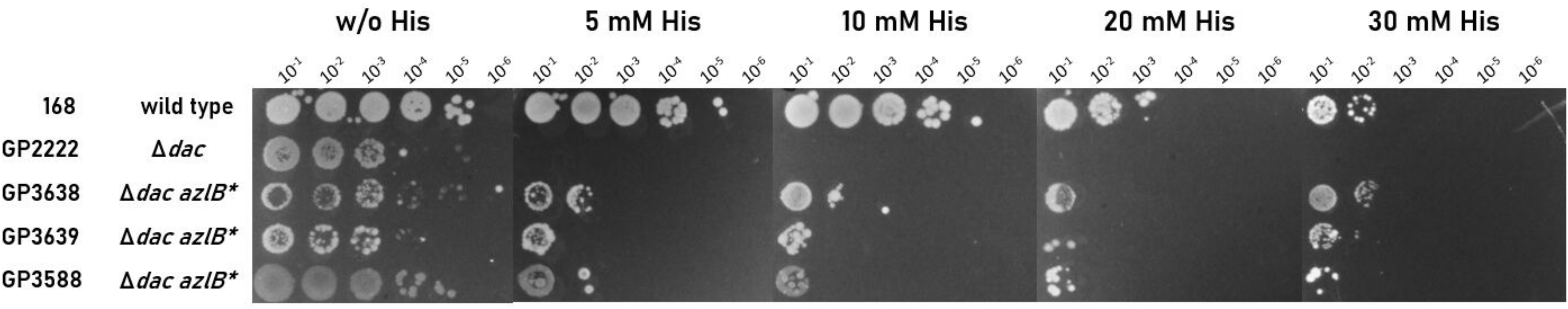
The isolated suppressors are resistant to histidine stress. Growth of *B. subtilis* suppressor mutants (GP3638, GP3639 and GP3588) in the presence of histidine. All suppressors carry different mutations in the *azlB* gene (see Table 2). Cells were harvested and washed, and the OD_600_ was adjusted to 1.0. Serial dilutions were added dropwise to MSSM minimal plates with the indicated histidine concentration. Plates were incubated at 42°C for 48 h.

As mentioned above, growth of the wild type strain 168 was inhibited above 20 mM histidine. Therefore, we tested the growth of our suppressor mutants in the absence and presence of histidine. While all mutants were viable at 5 mM histidine, they were still inhibited at a concentration of 30 mM (see Fig. 1). However, when suppressor mutants originally isolated at 15 mM histidine were transferred to 30 mM histidine, suppressor mutants appeared again. One of these mutants (GP3588) was subjected to whole genome sequencing. In coherence with our previously isolated mutants, we found a frameshift mutation in *azlB* highlighting the importance of *azlB* inactivation for the adaptation of the *B. subtilis* strain lacking c-di-AMP to the presence of histidine. Moreover, we found three additional mutations at a histidine concentration of 30 mM. Both the main potassium transporter KimA and the KtrD membrane subunit of the low-affinity potassium channel KtrCD (21, 27) were inactivated due to frameshift mutations. In addition, the high affinity glutamate transporter GltT (14, 28) carried a substitution of Thr-342 to a Pro residue. It is known that KtrCD is converted to a high-affinity potassium channel in the presence of glutamate (22) suggesting that glutamate as the product of histidine utilization causes activation of KtrCD. Moreover, small amounts of glutamate that are exported from the cell may be re-imported by GltT thus, again, contributing to the activation of KtrCD. This activation of KtrCD as well as the activity of KimA contribute to potassium toxicity that can only be bypassed by inactivation of the major potassium uptake systems.

Histidine toxicity in the *Δdac* mutant GP2222 might be caused by the formation of glutamate that triggers toxic glutamate accumulation, by toxicity of histidine due to its reactivity itself, or by a combination of both. The identification of *kimA* and *ktrD* mutants in the suppressor isolated at the elevated histidine concentration suggests that potassium toxicity really can become a problem for the bacteria. However, we never identified suppressor mutants that were affected in the histidine degradaton pathway thus avoiding the problem of intracellular glutamate formation. To test the role of histidine degradation for the acquisition of resistance to histidine, we deleted the *hutH* gene encoding histidase, the first gene of the catabolic pathway in a wild type and a *Δdac* mutant strain. The set of four isogenic strains was tested for growth on minimal medium in the absence of histidine and in the presence of 5 mM, 15 mM, 25 mM, and 35 mM histidine. While all strains grew well in the absence of histidine, growth was inhibited at histidine concentrations exceeding 15 mM and 5 mM for the wild type and the *Δdac* mutant, respectively. The inactivation of the histidine degradative pathway *(hutH* mutation) did not affect growth in either of the genetic backgrounds (data not shown). Thus, histidine seems to inhibit growth of *B. subtilis* by itself, as has also been observed for *E. coli* (29).

Taken together, our results demonstrate that the Lrp-type repressor protein AzlB plays a major role in the adaptation of *B. subtilis* lacking c-di-AMP to high levels of histidine. At even higher concentration of histidine, the degradation product glutamate induces the uptake of potassium, which is known to be toxic to strains that are unable to produce c-di-AMP (20, 21, 22).

### Suppressor mutants exhibit increased expression of the *azlBCD-brnQ* operon

To test the effect of the *azlB* mutations on the expression of the *azlBCD-brnQ* operon, we analyzed the transcripts of the operon by a Northern blot analysis. For this purpose, we cultivated the wild type strain 168, the *Δdac* mutant GP2222 and two suppressor mutants GP3638 (AzlB-Asn24 Ser) and GP3639 (frameshift in AzlB) in MSSM minimal medium, isolated the RNA, and performed Northern blot experiments using a riboprobe complementary to *azlC* to detect the specific mRNA(s). Based on the known transcript sizes of the *B. subtilis* glycolytic *gapA* operon (30), we estimated the sizes of the transcripts of the *azl* operon. As shown in Fig. 2A, expression of the operon could not be detected in the wild type strain 168 and in the *Δdac* mutant GP2222. Signals corresponding to transcripts of about 1,100, 1,500, 3,300 and 5,100 nucleotides were only visible in the two suppressor mutants carrying the *azlB* mutations. This result indicates that only inactive *azlB* allows expression of the *azl* operon and led to high expression levels. The presence of multiple transcripts suggests internal transcription signals and/or mRNA processing events.

**Fig. 2.**
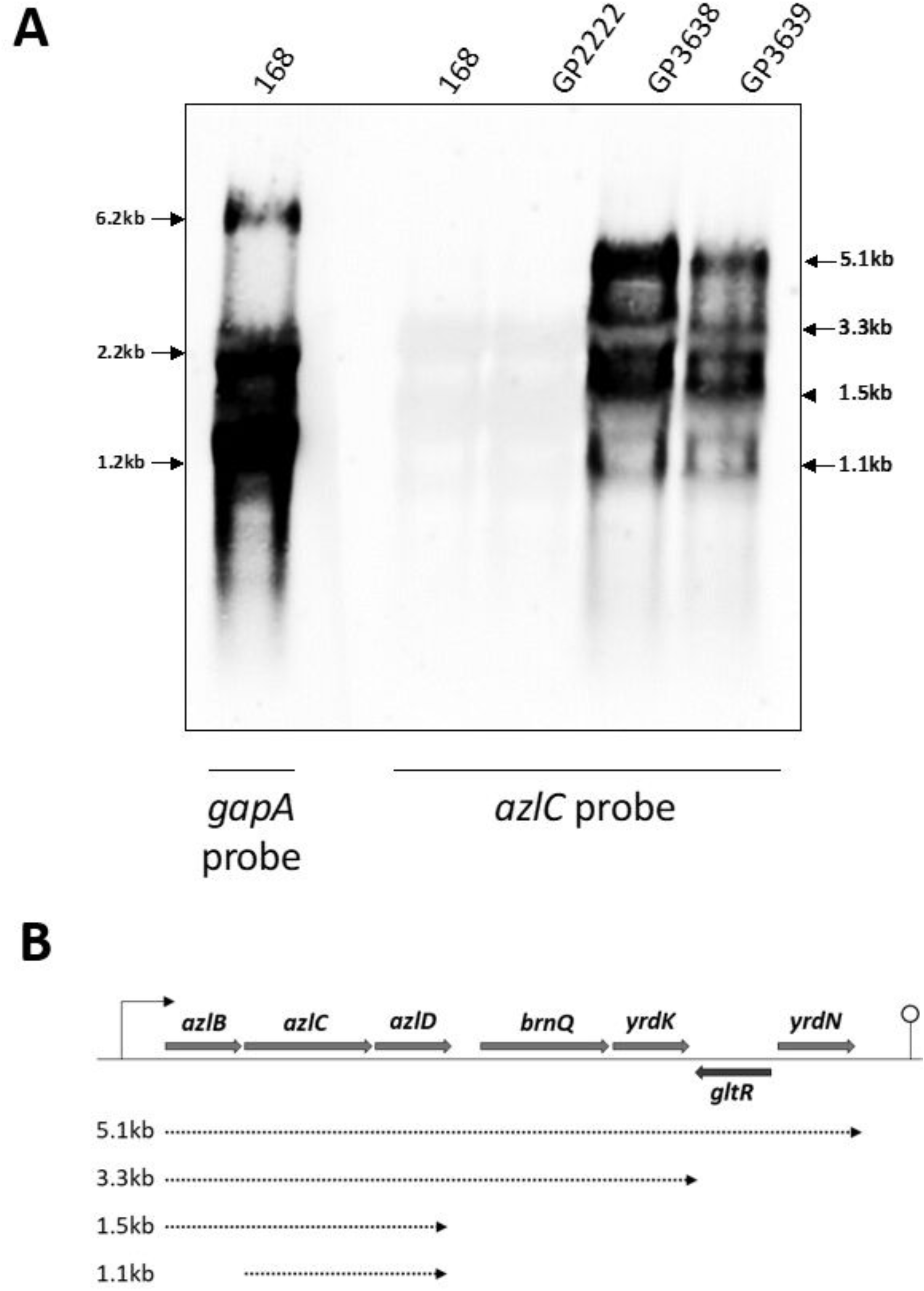
AzlCD is strongly overexpressed in the histidine suppressors. **A.** Northern Blot analysis to test the expression levels of the *azl* operon in the suppressor mutants GP3638 and GP3639. The RNA was isolated from MSSM minimal medium during exponential growth phase. The *gapA* probe was used together with wild type RNA as control to estimate band sizes and strength, as it is strongly expressed under normal conditions. **B.** Transcriptional organization of the *azl* operon. The length of the individual transcripts is indicated by the arrows.

So far, the inducer for the *azlBCD-brnQ* operon has not been identified. Since our results indicate that the operon is involved in the control of the histidine homeostasis, we wanted to test the activity of the *azlB* promoter under different conditions. For this purpose, we fused the *azlB* promoter region to a promoterless *lacZ* gene encoding β-galactosidase and integrated this *azlB-lacZ* fusion into the *B. subtilis* genome. According to a genome-wide transcriptome analysis (31), the promoter of the operon is located in front of the upstream *yrdF* gene. However, the same study indicated the presence of an mRNA upshift in front of the *azlB* gene. Therefore, we also constructed and tested an *yrdF-lacZ* fusion. The strains carrying the *azlB-lacZ* and *yrdF-lacZ* fusions were cultivated in C-Glc minimal medium in the absence or presence of different amino acids as potential inducers. As shown in Table 1, both the upstream regions of *yrdF* and *azlB* had only very minor promoter activity. As a control, we used the moderately expressed promoter of the *pgi* gene encoding phosphoglucose isomerase. This promoter yielded a ten-fold higher β-galactosidase activity as compared to the *yrdF* and *azlB* promoters. In addition, the activity of the *yrdF* and *azlB* promoters was not induced by any of the tested amino acids, including histidine. Therefore, we also tested casein hydrolysate, a mixture of amino acids. Again, no induction was observed for both promoters. However, deletion of the *azlB* gene resulted in an about seven-fold increase of the activity of the *azlB* promoter (see GP3614 vs. GP3612) whereas the *yrdF* promoter was not affected. Moreover, GltR, a LysR family transcription factor of so far unknown function, is encoded downstream of the *brnQ* gene (32). We therefore considered the possibility that GltR might play a role in the control of the *azl* operon. However, deletion of the *gltR* gene did not affect the activity of the *azlB* promoter.

**Table 1.** Activity of the *azlB* promoter

| Strain | Relevant genotype | Units of β-galactosidase per $\mu\text{g}$ of protein | | | | |
| --- | --- | --- | --- | --- | --- | --- |
|  |  | Addition to C-Glc minimal medium |  |  |  |  |
|  |  | - | Ile | Pro | His | CAA |
| GP314 | *pgi-lacZ* | $48 \pm 3$ | ND | ND | ND | ND |
| GP3612 | *azlB-lacZ* | $4 \pm 0.6$ | $3 \pm 0.7$ | $2 \pm 0.2$ | $4 \pm 0.6$ | $7 \pm 1.5$ |
| GP3614 | *azlB-lacZ \Delta azlB* | $26 \pm 5$ | $25 \pm 2$ | $31 \pm 4$ | $31 \pm 4$ | $49 \pm 7$ |
| GP3617 | *azlB-lacZ \Delta gltR* | $3 \pm 0.3$ | ND | ND | ND | $5 \pm 0.1$ |
| GP3611 | *yrdF-lacZ* | $3 \pm 0.7$ | $4 \pm 1$ | $2 \pm 0.3$ | $5 \pm 2$ | ND |
| GP3613 | *yrdF-lacZ \Delta azlB* | $3 \pm 0.3$ | $3 \pm 0.6$ | $3 \pm 0.1$ | $3 \pm 0.4$ | ND |

Taken together, our data confirm that AzlB is the transcriptional repressor of the *azl* operon. The *azlB* gene is the first gene of the operon (see Fig. 2B). Moreover, our results demonstrate that the transcriptional regulation by AzlB is not affected by any individual amino acid or a mixure of them, even though the operon encodes exporters and importers for amino acids. Only the loss of a functional AzlB repressor allows the expression of the *azl* operon (see Discussion).

### Resistance to histidine depends on the AzlCD amino acid exporter

So far, we have established that the suppressor mutants have mutations in AzlB that increase expression of the *azl* operon, which confers resistance to histidine. In addition to the promoter-proximal repressor gene *azlB*, this operon encodes the AzlC and AzlD subunits of a bipartite amino acid exporter and the branched-chain amino acid transporter BrnQ as well as the YrdK protein of unknown function and the putative 4-oxalocrotonate tautomerase YrdN. Since overexpression of AzlCD was also responsible for the resistance of *B. subtilis* to azaleucine (26), it seemed most plausible that this transporter is also involved in histidine resistance. To test this hypothesis, we constructed two sets of isogenic strains that differed in the *azl* operon in the background of the wildtype 168 and in the background of the *Δdac* mutant GP2222. First, we compared growth of the wild type, the *azlB* mutant GP3600, and the *azlBCD* mutant GP3601. As shown in Fig. 3A, the wild type strain was sensitive to the presence of 15 mM histidine in the medium, whereas the isogenic *azlB* mutant that exhibits overexpression of AzlCD was resistant. However, the additional deletion of the *azlCD* genes in GP3601 restored the sensitivity to histidine, indicating that the increased expression of the AzlCD amino acid exporter is responsible for the acquired resistance to histidine. Similar results were obtained for the set of strains that are unable to synthesize c-di-AMP (Δ*dac*, Fig. 3B). Again, the strain lacking AzlB (GP3607) was resistant to high levels of histidine (20 mM), whereas the strain lacking the amino acid exporter AzlCD in addition to AzlB (GP3606) was as sensitive as the *Δdac* mutant (GP2222) even at 5 mM histidine. Ectopic expression of the *azlCD* genes under the control of the constitutive *degQ36* promoter (33) in strain GP3642 that lacks the endogenous *azlBCD* operon partially restored the resistance to histidine up to a concentration of 5 mM. In contrast, expression of the AzlC component of the bipartite exporter alone had no effect (Fig. 3B, GP3643). Taken together, these data strongly suggest that the overexpression of the two-component amino acid exporter AzlCD as a result of the inactivation of AzlB is required for the resistance of *B. subtilis* to histidine.

**Fig. 3.**
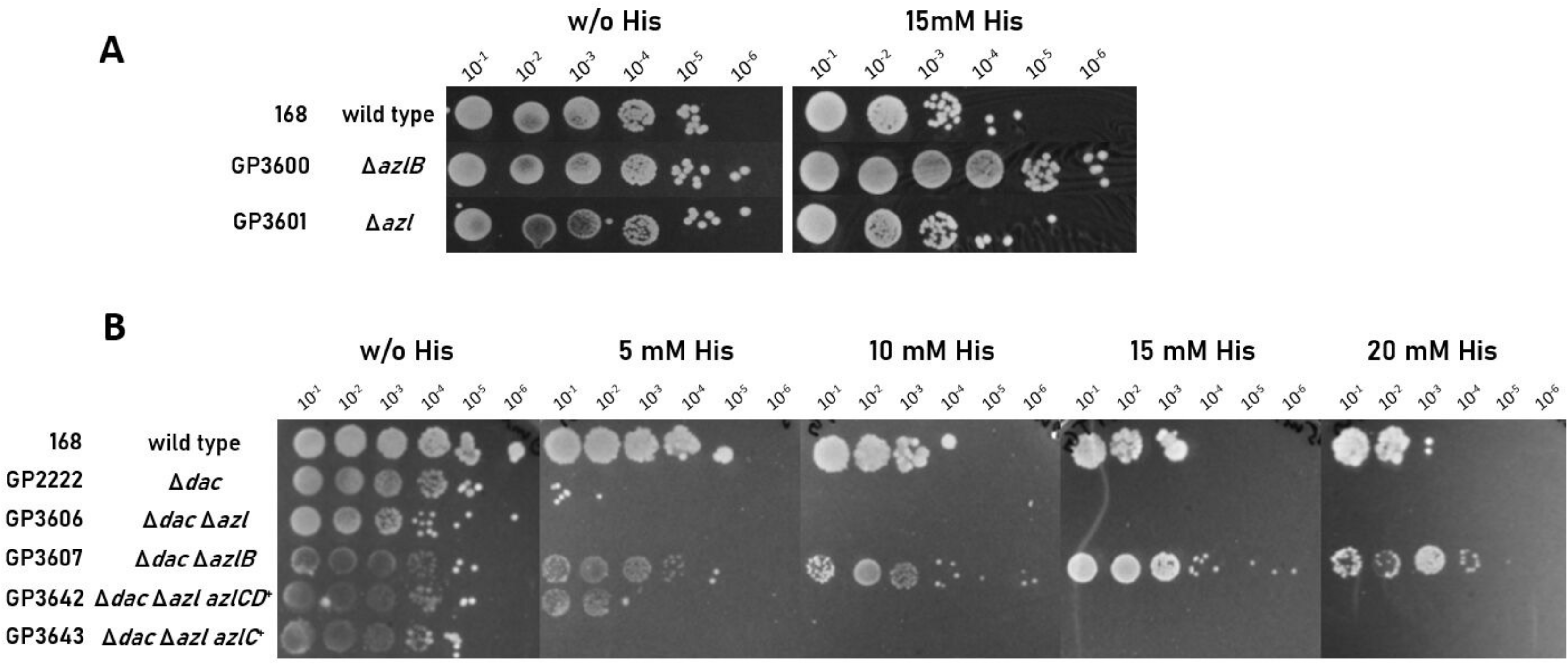
The *azlB* mutation confers resistance to histidine stress. **A.** Sensitivity of wild type *B. subtilis* (168) and the *ΔazlB* (GP3600) and *ΔazlBCD* (GP3601) mutants to histidine. Cells were grown in MSSM minimal medium to an OD_600_ of 1.0 and then diluted 10-fold to create dilutions ranging from 10^-1^ to 10^-6^. The dilution series was dropped onto MSSM plates without and with (15 mM) histidine. The plates were incubated at 37°C for 48 h. **B.** Growth of *Δdac Δazl* (GP3606), *Δdac ΔazlB* (GP3607) and *Δdac Δazl* complemented with *azlC* (GP3643) and *azlCD* (GP3642) respectively. *Δazl* indicates a deletion of the *azlBCD* genes. Cells were grown as described above. The plates were incubated at 42°C for 48 h.

### Overexpression of AzlCD results in enhanced histidine export

AzlCD has previously been identified as an exporter for 4-azaleucine and was hypothesized to be an exporter for other branched chain amino acids (26). Our data suggest that the complex might also export histidine thus contributing to histidine resistance upon overexpression. To test this idea, we determined the relative intra- and extracellular histidine concentrations in the wild type strain 168, as well as in the isogenic *azlB, azlBCD*, and *azlCD* deletion mutants GP3600, GP3601, and GP3622, respectively, during growth in MSSM minimal medium. In this condition, *de novo* histidine biosynthesis is active, because MSSM minimal medium does not contain amino acids. Compared to the wild type, intracellular histidine levels decreased in the *azlB* mutant GP3600, thus confirming that higher AzlCD levels in this strain led to histidine export (Fig. 4A). Mutants lacking the amino acid exporter AzlCD had wild type-like histidine levels (Fig. 4A). In contrast, the extracellular histidine concentration was threefold higher in the *azlB* mutant whereas the strains lacking AzlCD have extracellular histidine levels that were comparable to the wild type strain (Fig 4 B). These results demonstrate that AzlCD which is overexpressed as a result of the *azlB* mutation, is involved in the control of histidine homeostasis. While the loss of AzlCD has no effect, which corresponds to the lack of expression in the wild type strain, its overexpression results in reduced and increased intra- and extracellular histidine levels, respectively. This suggests that AzlCD is an active histidine exporter.

**Fig. 4.**
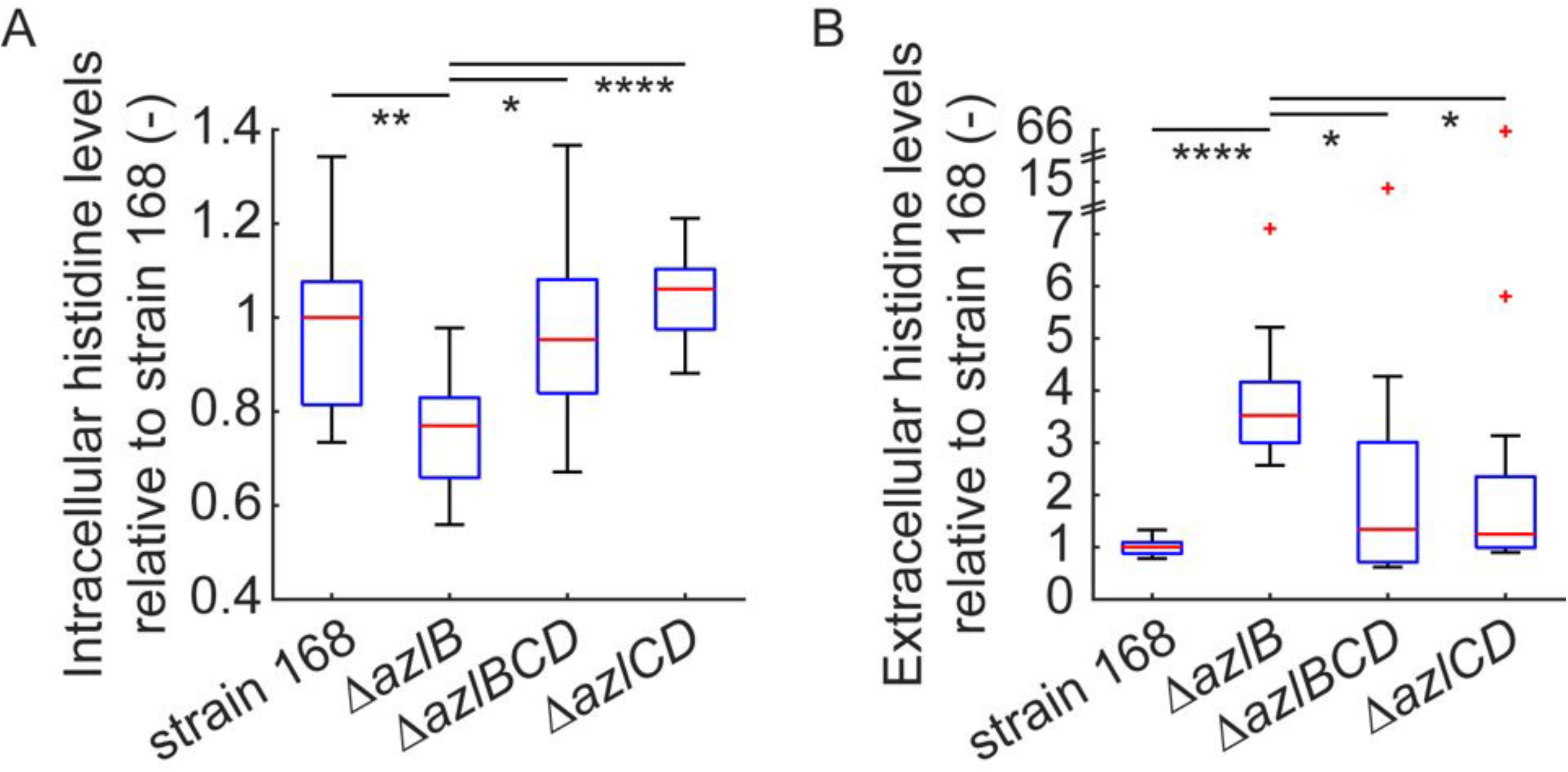
AzlCD is a histidine exporter in *B. subtilis*. Box-whisker plot of the intracellular (A) and extracellular (B) histidine levels of *B. subtilis ΔazlB, ΔazlBCD*, and *ΔazlCD* mutants relative to the wild type strain 168. The red lines indicate the median values of 12 biological replicates. The upper box edges show the 75^th^ percentiles, the lower edges the 25^th^ percentiles. The whiskers indicate the furthest points that are not considered outliers. The red crosses indicate outliers. Differences between indicated pairs of strains were tested for significance using a Wilcoxon rank sum test at a significance level α of 0.05. p-values < 0.05 were considered statistically significant. The stars indicate the orders of magnitude of the p-values: * = p < 0.05; ** = p < 0.01; *** = p < 0.001; **** = p < 0.0001.

## DISCUSSION

The results presented in this study demonstrate that histidine inhibits growth of *B. subtilis* as has already been shown for serine or threonine (10, 34, 35). Amino acid toxicity is often enhanced if *B. subtilis* is unable to produce the essential second messenger nucleotide c-di-AMP due to the activation of the potassium channel KtrCD by glutamate, the degradation product of many amino acids (14, 22). This work shows that the increased sensitivity of a strain lacking c-di-AMP to amino acids is also valid for histidine.

Typically, suppressor screens using toxic amino acids, amino acids analogs or related anti-metabolites result in the identification of transporters, which have been inactivated in the suppressor mutants (9, 10, 11, 12, 13, 14, 15). While this is the predominant type of suppressor mutations, resistance to toxic amino acids and related molecules can also be achieved by the activation of degradation pathways (10, 14), by the activation of export mechanisms (14, 26), or by modifying the target protein/ pathway in a way that it becomes insensitive to the presence of the otherwise toxic molecule. This was observed for glyphosate resistance in *Salmonella typhimurium*, which can be achieved by mutations that render the target enzyme 5-enolpyruvyl-shikimate-3-phosphate (EPSP) synthase insensitive to inhibition (18) as well as for serine toxicity in *B. subtilis*, which could be overcome by increased expression of the genes encoding the threonine biosynthetic pathway (10). Studies about histidine toxicity in *E. coli* revealed that the amino acid enhances oxidative DNA damage (29). Thus one might also expect suppressor mutations that prevent DNA damage. The exclusive isolation of *azlB* mutations that activate the expression of the AzlCD amino acid exporter suggests that all other mechanisms of suppression are beyond reach for the *B. subtilis* cell.

The fact that we were unable to isolate a single suppressor mutant that had lost histidine uptake strongly suggests that *B. subtilis* possesses multiple histidine transporters. So far, only the HutM histidine transporter has been identified based on its similarity to known histidine transporters (25). However, the expression of the *hutM* gene in the *hut* operon for histidine utilization as well as the induction of its expression by histidine suggest that HutM is a high-affinity transporter that is probably not involved in histidine uptake under standard conditions. It is thus tempting to speculate that the genome of *B. subtilis* encodes one or more low-affinity transporters for histidine. Indeed, *B. subtilis* encodes several homologs of the *Pseudomonas putida* histidine transporter HutT (36). These transporters all belong to the amino acid-polyamine-organocation (APC) superfamily of amino acid transporters. Four of them (AapA, AlaP, YbxG, and YdgF) share more than 40% sequence identity wih *P. putida* HutT, suggesting that these proteins have the same biological activity. Thus, the presence of multiple histidine uptake systems would prevent the rapid simultaneous inactivation of all these systems in suppressor mutants thus explaining that no transporter mutants were isolated.

Our data clearly demonstrate that the bipartite amino acid transporter AzlCD exports not only the leucine analog 4-azaleucine (16), but also histidine. Corresponding bipartite systems that mediate the export of branched-chain amino acids have also been identified in *E. coli* and *Corynebacterium glutamicum* (37, 38). These exporters are members of the LIV-E class of transport proteins (38, 39). As in *B. subtilis*, these systems consist of a large (corresponding to AzlC) and a small subunit (corresponding to AzlD). While proteins homologous to AzlC are abundant in a wide range of bacteria, including most Actinobacteria and Firmicutes as well as many Proteobacteria, AzlD is conserved only in few bacteria. The other bacteria that possess a homolog of AzlC obviously have alternative small subunits. This is the case in *E. coli*, where the small YgaH subunit of the YgaZ/YgaH valine exporter is not similar to its counterparts in *B. subtilis* and *C. glutamicum*. We have also considered the possibility that the large subunit AzlC might be sufficient for histidine export; however, this is not the case (see Fig. 3B).

It is interesting to note that the AzlCD amino acid exporter is able to export multiple amino acids. Substrate promiscuity is a common feature in amino acid transport. In *B. subtilis*, the low affinity transporter AimA is the major transporter for glutamate and serine (10, 14). Similarly, the BcaP permease transports branched-chain amino acids, threonine and serine (9, 10, 11) and the GltT protein is involved in the uptake of aspartate, glutamate, and the antimetabolite glyphosate (14, 15, 28). Thus, AzlCD is another example for the weak substrate specificity of amino acid transporters. It is tempting to speculate that AzlCD might be involved in the export of even other amino acids and related metabolites in *B. subtilis*.

Based on the chemical properties of each amino acid, it may be generally toxic, or only under specific conditions. Therefore, cells often have efficient degradation pathways to remove toxic compounds. This is the case for glutamate which is degraded by the glutamate dehydrogenases GudB or RocG (7, 17). However, other amino acids become toxic only at very high concentrations or in very particular mutant backgrounds. This is the case for histidine which is toxic only at high concentrations for the *B. subtilis* wild type strain but already at low concentrations in a strain unable to form c-di-AMP. Similarly, the presence of amino acid analogs such as 4-azaleucine might be a rather exceptional event in natural environments. Still, *B. subtilis* is equipped to meet this challenge using the amino acid exporter AzlCD. Based on a global transcriptome analysis, the *azl* operon is barely expressed under a wide range of conditions, and no conditions that results in induction of the operon could be detected (26, 31). Similarly, the putative arginine and lysine exporter YisU is not expressed under any of 104 studied conditions (31). The observation that the expression of the *azl* operon in the presence of toxic concentrations of histidine or 4-azaleucine is obviously not sufficient to provide resistance against these amino acids already suggested that none of these compounds acts as a molecular inducer for the *azl* operon. In agreement with previous results (26), we observed substantial expression of the operon only if the *azlB* gene encoding the repressor of the operon was deleted or inactivated due to the suppressor mutations. Even the presence of a mixture of amino acids derived from casamino acids did not result in the induction of the operon. As the functions of the operon seem to be related to amino acid export (AzlCD) and uptake (BrnQ), regulation by amino acid availability seemed to be most likely. However, the results from prior global and operon-specific transcription studies as well as our data suggest that the activity of AzlB is not controlled by amino acids even though the protein belongs to Lrp family of leucine-responsive regulatory proteins (40). It is tempting to speculate that AzlB has lost the ability to interact with amino acid-related effector molecules, but that expression of the operon can rapidly be activated by the acquisition of mutations that inactivate AzlB. Alternatively, AzlB might respond to a yet unknown signal and then allow induction of the operon. The mutational inactivation of a normally silent operon has also been described for the cryptic *E. coli bgl* operon for the utilization of β-glucosides which requires insertion of the mobile element IS5 in the promoter region to get expressed (41).

Due to its strongly increased sensitivity to several amino acids, the *B. subtilis* mutant lacking c-di-AMP is an excellent tool to study mechanisms of amino acid homeostasis, and to identify uptake and export systems. This endeavour is required as the details of amino acid transports are one of the few areas, which has several gaps of knowledge in the research on *B. subtilis* (3). We anticipate that the further use of the c-di-AMP lacking mutant will continue to help filling these remaining gaps.

## MATERIALS AND METHODS

### Strains, media and growth conditions

*E. coli* DH5α (42) was used for cloning. All *B. subtilis* strains used in this study are derivatives of the laboratory strain 168. They are listed in Table 2. *B. subtilis* and *E. coli* were grown in Luria-Bertani (LB) or in sporulation (SP) medium (42, 43). For growth assays, *B. subtilis* was cultivated in MSSM medium (21). MSSM is a modified SM medium in which KH_2_PO_4_ was replaced by NaH_2_PO_4_ and KCl was added as indicated (21). The media were supplemented with ampicillin (100 μg/ml), kanamycin (10 μg/ml), chloramphenicol (5 μg/ml), spectinomycin (150 μg/ml), tetracycline (12.5 μg/ml) or erythromycin and lincomycin (2 and 25 μg/ml, respectively) if required.

**Table 2.**
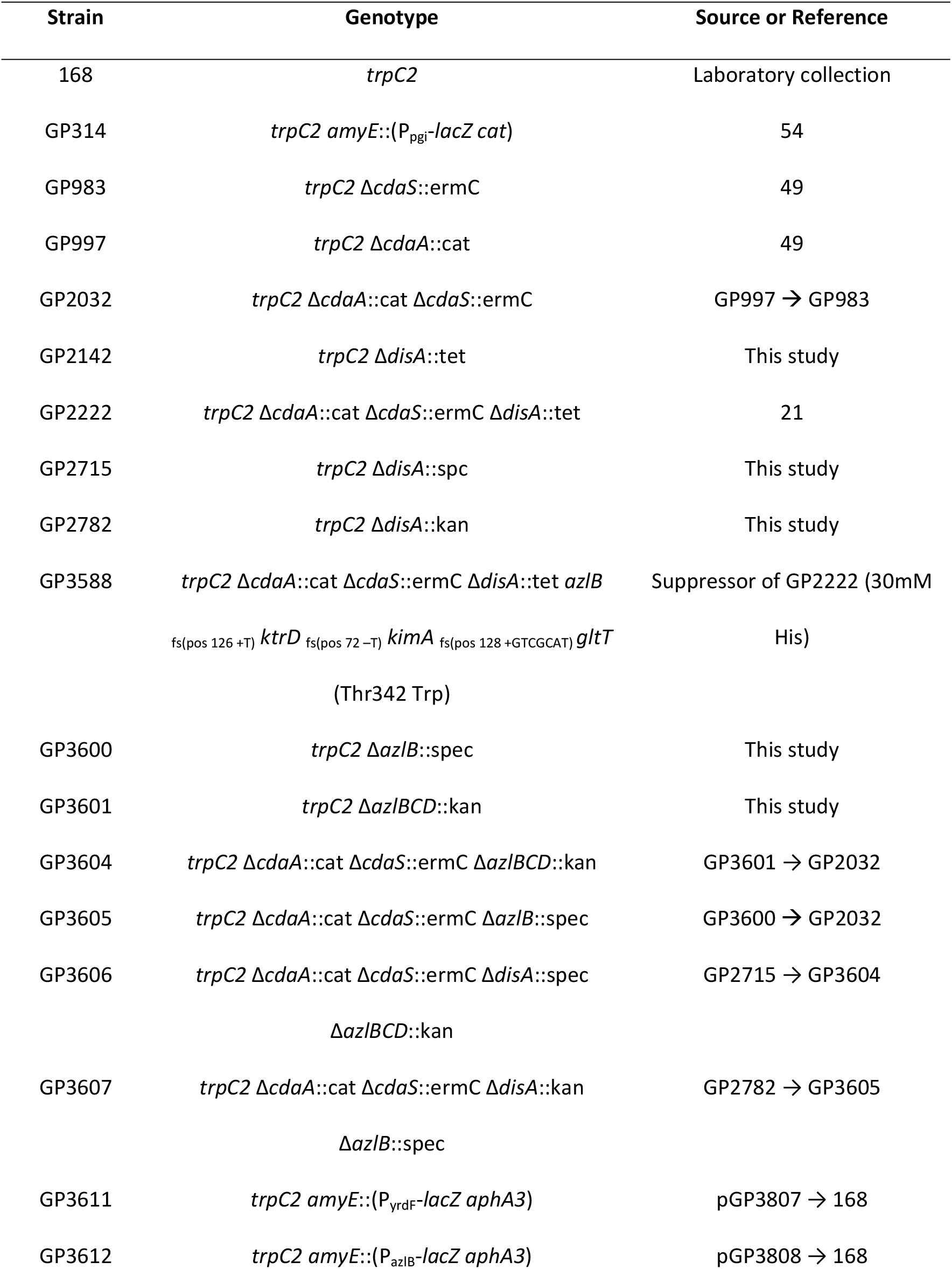
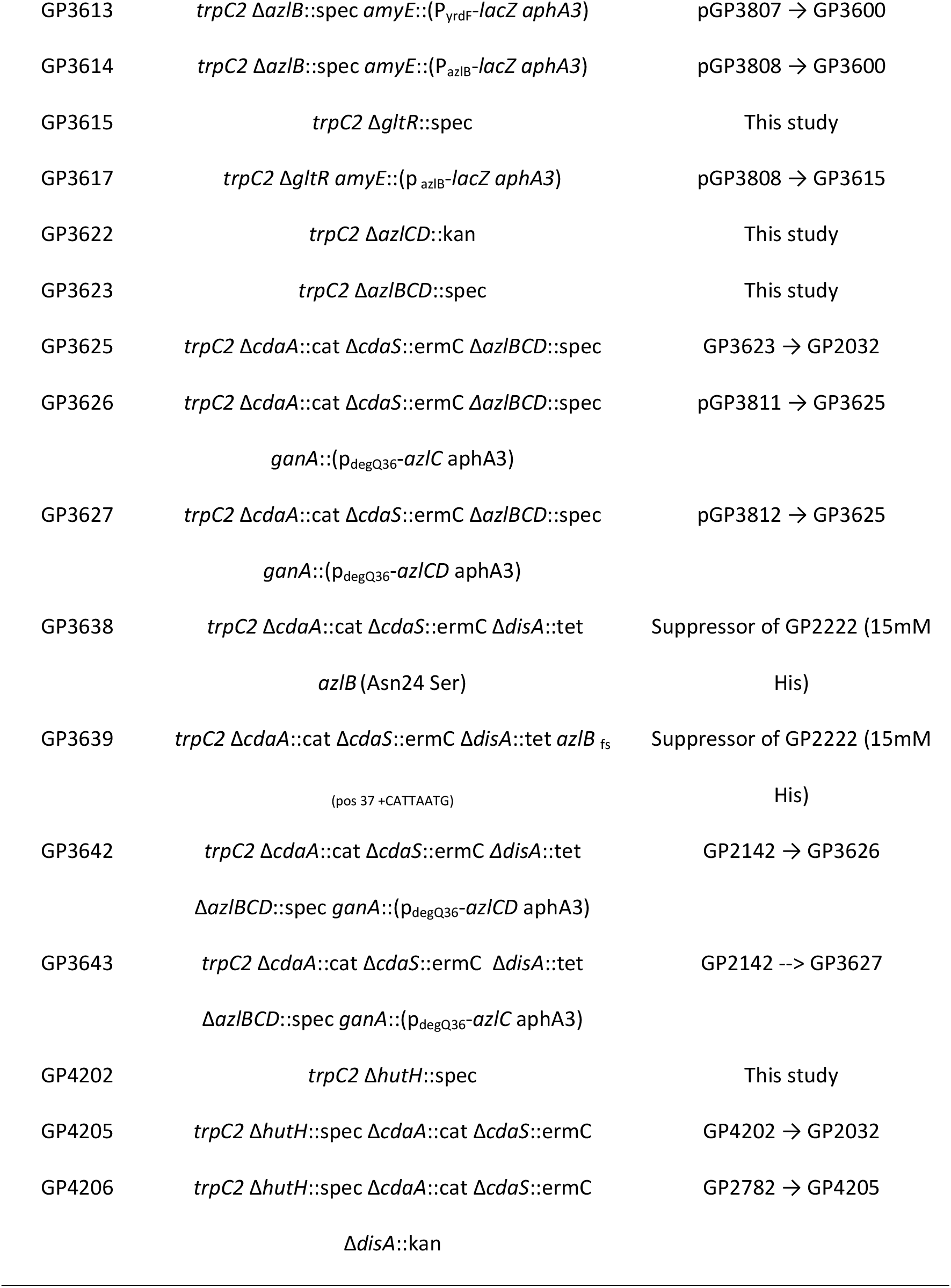
*B. subtilis* strains used in this study

### DNA manipulation and transformation

All commercially available restriction enzymes, T4 DNA ligase and DNA polymerases were used as recommended by the manufacturers. DNA fragments were purified using the QIAquick PCR Purification Kit (Qiagen, Hilden, Germany). DNA sequences were determined by the dideoxy chain termination method (42). Standard procedures were used to transform *E. coli* (42), and transformants were selected on LB plates containing ampicillin (100 μg/ml). *B. subtilis* was transformed with plasmid or chromosomal DNA according to the two-step protocol described previously (43). Transformants were selected on SP plates containing chloramphenicol (Cm 5 μg/ml), kanamycin (Km 10 μg/ml), spectinomycin (Spc 150 μg/ml), tetracycline (Tet 12,5 μg/ml) or erythromycin plus lincomycin (Em 2 μg/ml and Lin 25 μg/ml).

### Genome sequencing

To identify the mutations in the suppressor mutant strains GP3588, GP3638, and GP3639 (see Table 2), the genomic DNA was subjected to whole-genome sequencing. Concentration and purity of the isolated DNA was first checked with a Nanodrop ND-1000 (PeqLab Erlangen, Germany), and the precise concentration was determined using the Qubit® dsDNA HS Assay Kit as recommended by the manufacturer (Life Technologies GmbH, Darmstadt, Germany). Illumina shotgun libraries were prepared using the Nextera XT DNA Sample Preparation Kit and subsequently sequenced on a MiSeq system with the reagent kit v3 with 600 cycles (Illumina, San Diego, CA, USA) as recommended by the manufacturer. The reads were mapped on the reference genome of *B. subtilis* 168 (GenBank accession number: NC_000964) (44). Mapping of the reads was performed using the Geneious software package (Biomatters Ltd., New Zealand) (45). Frequently occurring hitchhiker mutations (46) and silent mutations were omitted from the screen. The resulting genome sequences were compared to that of our in-house wild type strain. Single nucleotide polymorphisms were considered as significant when the total coverage depth exceeded 25 reads with a variant frequency of ≥90%. All identified mutations were verified by PCR amplification and Sanger sequencing.

### Construction of mutant strains by allelic replacement

Deletion of the *azlB, azlBCD, azlCD, disA, gltR*, and *hutH* genes was achieved by transformation of *B. subtilis* 168 with PCR product constructed using oligonucleotides to amplify DNA fragments flanking the target genes and an appropriate intervening resistance cassette as described previously (47). The integrity of the regions flanking the integrated resistance cassette was verified by sequencing PCR products of about 1,100 bp amplified from chromosomal DNA of the resulting mutant strains. In the case of the *azlB, azlCD*, and *azlBCD* deletions, the cassette carrying the resistance gene lacked a transcription terminator to ensure the expression of the downstream genes.

### Phenotypic analysis

In *B. subtilis*, amylase activity was detected after growth on plates containing nutrient broth (7.5 g/l), 17 g Bacto agar/l (Difco) and 5 g hydrolyzed starch/l (Connaught). Starch degradation was detected by sublimating iodine onto the plates.

Quantitative studies of *lacZ* expression in *B. subtilis* were performed as follows: cells were grown in MSSM medium supplemented with KCl at different concentrations as indicated. Cells were harvested at OD600 of 0.5 to 0.8. β-Galactosidase specific activities were determined with cell extracts obtained by lysozyme treatment as described previously (43). One unit of β-galactosidase is defined as the amount of enzyme which produces 1 nmol of o-nitrophenol per min at 28° C.

To assay growth of *B. subtilis* mutants at different histidine concentrations, a drop dilution assay was performed. Briefly, precultures in MSSM medium at the indicated histidine concentration were washed three times, resuspended to an OD_600_ of 1.0 in MSSM basal salts solution. Dilution series were then pipetted onto MSSM plates containing the desired histidine concentration.

### Plasmid constructions

Plasmid pAC7 (48) was used to construct translational fusions of the potential *yrdF* and *azlB* promoter regions to the promoterless *lacZ* gene. For this purpose, the promoter regions were amplified using oligonucleotides that attached EcoRI and BamHI restriction to the ends of the products. The fragments were cloned between the EcoRI and BamHI sites of pAC7. The resulting plasmids were pGP3807 and pGP3808 for *yrdF* and *azlB*, respectively.

To allow ectopic expression of the *azlC* and *azlCD* genes, we constructed the plasmids pGP3811 and pGP3812, respectively. The corresponding genes were amplified using oligonucleotides that added BamHI and PstI sites to the ends of the fragments and cloned into the integrative expression vector pGP1460 (49) linearized with the same enzymes.

### Northern blot analysis

The strains *B. subtilis* 168 (wild type) and GP2222 (Δ*dac* mutant) as well as the suppressor mutants GP3638 and GP3639 were grown in MSSM minimal medium and harvested in the late logarithmic phase. The preparation of total RNA and Northern blot analysis were carried out as described previously (50, 51). Digoxigenin (DIG) RNA probes were obtained by *in vitro* transcription with T7 RNA polymerase (Roche Diagnostics) using PCR-generated DNA fragments as templates. The reverse primer contained a T7 RNA polymerase recognition sequence. *In vitro* RNA labelling, hybridization and signal detection were carried out according to the instructions of the manufacturer (DIG RNA labelling kit and detection chemicals; Roche Diagnostics).

### Determination of intra- and extracellular histidine pools

For the determination of histidine levels of *B. subtilis*, cells were cultivated in MSSM minimal medium until exponential growth phase (OD_600_ of 0.4). For the extraction of intracellular metabolites, 4 ml of each culture were harvested by filtration (52). Histidine levels were then determined as described previously (53) using ^13^C labelled histidine from an *E. coli* extract as internal standard. Briefly, an Agilent 1290 Infinity II UHPLC system (Agilent Technologies) was used for liquid chromatography. The column was an Acquity BEH Amide 30 x 2.1 mm with 1.7 μm particle size (Waters GmbH). The temperature of the column oven was 30°C, and the injection volume was 3 μl. LC solvent A was: water with 10 mM ammonium formate and 0.1 % formic acid (v/v), and LC solvent B was: acetonitrile with 0.1 % formic acid (v/v). The gradient was: 0 min 90% B; 1.3 min 40 % B; 1.5 min 40 % B; 1.7 min 90 % B; 2 min 90 % B; 2.75 min 90% B. The flow rate was 0.4 ml min^-1^. From minute 1 to 2, the sample was injected to the MS. An Agilent 6495 triple quadrupole mass spectrometer (Agilent Technologies) was used for mass spectrometry. Source gas temperature was set to 200°C, with 14 l min^-1^ drying gas and a nebulizer pressure of 24 psi. Sheath gas temperature was set to 300°C and flow to 11 l min^-1^. Electrospray nozzle and capillary voltages were set to 500 and 2500 V, respectively. Isotope-ratio mass spectrometry with ^13^C internal standard was used to obtain relative data. Fully ^12^C- and ^13^C-labelled histidine was measured by multiple reaction monitoring in positive ionization mode using a collision energy of 13 eV. Precursor ion masses were 156 Da and 162 Da, product ion masses 110 Da and 115 Da for ^12^C- and ^13^C-histidine, respectively. Ratios between ^12^C- and ^13^C-labelled histidine were normalized to the ODs and the median ratio of the control strain 168.

## ACKNOWLEDGEMENTS

This work was supported by a grant of the Deutsche Forschungsgemeinschaft (DFG) within the Priority Program SPP1879 (Stu 214-16) (to J.S.). H.L. and T.S. acknowledge funding from the Cluster of Excellence EXC 2124 from the Deutsche Forschungsgemeinschaft. The funders had no role in study design, data collection, analysis and interpretation, decision to submit the work for publication, or preparation of the manuscript. Anja Poehlein und Rolf Daniel are acknowledged for the genome sequencing.

## Author contributions

Design of the study: J.M. and J.S. Experimental work: J.M., T.S., B.M.H., and K.S. Data analysis: J.M., T.S., H.L. and J.S. Wrote the paper: J.M. and J.S.

## REFERENCES

1. De Lorenzo V, Sekowska A, Danchin A. 2015. Chemical reactivity drives spatiotemporal organization of bacterial metabolism. FEMS Microbiol Rev 39:96–119.

2. Commichau FM, Pietack FM, Stülke J. 2013. Essential genes in *Bacillus subtilis:* a re-evaluation after ten years. Mol Biosyst 9:1068–1075.

3. Reuß DR, Commichau FM, Gundlach J, Zhu B, Stülke J. 2016. The blueprint of a minimal cell: MiniBacillus. Microbiol Mol Biol Rev 80:955–987.

4. Aguilar Suárez R, Stülke J, van Dijl JM. 2019. ess is more: toward a genome-reduced *Bacillus* cell factory for “difficult proteins”. ACS Synth Biol 8:99–108.

5. Van Tilburg AY, van Heel AJ, Stülke J, de Kok NAW, Rueff AS, Kuipers OP. 2020. MiniBacillus PG10 as a convenient and effective production host for lantibiotics. ACS Synth Biol 9:1833–1842.

6. Michalik S, Reder A, Richts B, Faßhauer P, Mäder U, Pedreira T, Poehlein A, van Heel AJ, van Tilburg AY, Altenbuchner J, Klewing A, Reuß DR, Daniel R, Commichau FM, Kuipers OP, Hamoen LW, Völker U, Stülke J. 2021. The *Bacillus subtilis* minimal genome compendium. ACS Synth Biol 10:2767–2771.

7. Commichau FM, Gunka K, Landmann JJ, Stülke J. 2008. Glutamate metabolism in *Bacillus subtilis:*gene expression and enzyme activities evolved to avoid futile cycles and to allow rapid responses to perturbations of the system. J Bacteriol 190:3557–3564.

8. Gunka K, Tholen S, Gerwig J, Herzberg C, Stülke J, Commichau FM. 2012. A high-frequency mutation in *Bacillus subtilis:* requirements for the decryptification of the *gudB* glutamate dehydrogenase gene. J Bacteriol 194:1036–1044.

9. Belitsky BR. 2015. Role of branched-chain amino acid transport in *Bacillus subtilis* CodY activity. J Bacteriol 197:1330–1338.

10. Klewing A, Koo BM, Krüger L, Poehlein A, Reuß D, Daniel R, Gross CA, Stülke J. 2020. Resistance to serine in *Bacillus subtilis:* identification of the serine transporter YbeC and of a metabolic network that links serine and threonine metabolism. Environ Microbiol. 22:3937–3949.

11. Commichau FM, Alzinger A, Sande R, Bretzel W, Reuß DR, Dormeyer M, Chevreux B, Schuldes J, Daniel R, Akeroyd M, Wyss M, Hohmann HP, Prágai Z. 2015. Engineering *Bacillus subtilis* for the conversion of the antimetabolite 4-hydroxy-L-threonine to pyridoxine. Metab Eng 29:196–207.

12. Zaprasis A, Hoffmann T, Stannek L, Gunka K, Commichau FM, Bremer E. 2014. The γ-aminobutyrate permease GabP serves as the third proline transporter of *Bacillus subtilis*. J Bacteriol 196:515–526.

13. Sidiq KR, Chow MW, Zhao Z, Daniel RA. 2021. Alanine metabolism in *Bacillus subtilis*. Mol Microbiol 115:739–757.

14. Krüger L, Herzberg C, Rath H, Pedreira P, Ischebeck T, Poehlein A, Gundlach J, Daniel R, Völker U, Mäder U, Stülke J. 2021. Essentiality of c-di-AMP in *Bacillus subtilis:* bypassing mutations converge in potassium and glutamate homeostasis. PLoS Genet 17:e1009092.

15. Wicke D, Schulz LM, Lentes S, Scholz P, Poehlein A, Gibhardt J, Daniel R, Ischebeck T, Commichau FM. 2019. Identification oft he first glyphosate transporter by genomic adaptation. Environ Microbiol. 21:1287–1305.

16. Ward JB, Zahler SA. 1973. Regulation of leucine biosynthesis in *Bacillus subtilis*. J Bacteriol 116:727–735.

17. Belitsky BR, Sonenshein AL. 1998. Role and regulation of Bacillus subtilis glutamate dehydrogenase genes. J Bacteriol 180:6298–6305.

18. Comai L, Sen LC, Stalker DM. 1983. An altered *aroA* gene product confers resistance to the herbicide glyphosate. Science 221:370–371.

19. Stülke J, Krüger L. 2020. Cyclic di-AMP signaling in bacteria. Annu Rev Microbiol 74:159–179.

20. Gundlach J, Krüger L, Herzberg C, Turdiev A, Poehlein A, Tascón I, Weiß M, Hertel D, Daniel R, Hänelt I, Lee VT, Stülke J. 2019. Sustained sensing in potassium homeostasis: Cyclic di-AMP controls potassium uptake by KimA at the levels of expression and activity. J Biol Chem 294:9605–9614.

21. Gundlach J, Herzberg C, Kaever V, Gunka K, Hoffmann T, Weiß M, Gibhardt J, Thürmer A, Hertel D, Daniel R, Bremer E, Commichau FM, Stülke J. 2017. Control of potassium homeostasis is an essential function of the second messenger cyclic di-AMP in *Bacillus subtilis*. Sci Signal 10: eaal3011.

22. Krüger L, Herzberg C, Warneke R, Poehlein A, Stautz J, Weiß M, Daniel R, Hänelt I, Stülke J. 2020. Two ways to convert a low affinity potassium channel to high affinity: control of *Bacillus subtilis* KtrDC by glutamate. J Bacteriol 202:e00138–20.

23. Pedreira T, Elfmann C, Stülke J. 2022. The current state of *SubtiWiki*, the database for the model organism *Bacillus subtilis*. Nucleic Acids Res 50:D875–D882.

24. Wray LV, Fisher SH. 1994. Analysis of *Bacillus subtilis hut* operon expression indicates that histidine-dependent induction is mediated primarily by transcriptional antitermination and that amino acid repression is mediated by two mechanisms: regulation of transcription initiation and inhibition of histidine transport. J Bacteriol 176:5466–5473.

25. Bender RA. 2012. Regulation of the histidine utilization (*hut*) system in bacteria. Microbiol Mol Biol Rev 76:565–584.

26. Belitsky BR, Gustafsson MCU, Sonenshein AL, von Wachenfeldt C. 1997. An *lrp-*like gene of *Bacillus subtilis* involved in branched-chain amino acid transport. J Bacteriol 176:5466–5473.

27. Holtmann G, Bakker EP, Uozumi N, Bremer E. 2003. KtrAB and KtrCD: two K^+^ uptake systems in *Bacillus subtilis* and their role in adaptation to hypertonicity. J Bacteriol 185:1289–1298.

28. Zaprasis A, Bleisteiner M, Kerres A, Hoffmann T, Bremer E. 2015. Uptake of amino acids and their metabolic conversion into the compatible solute proline confers osmoprotection to *Bacillus subtilis*. Appl Environ Microbiol 81:250–259.

29. Nagao T, Nakayama-Imaohji H, Elahi M, Tada A, Toyonaga E, Yamasaki H, Okazaki K, Miyoshi H, Tsuchiya K, Kuwahara T. 2018. L-histidine augments the oxidative damage against Gram-negative bacteria by hydrogen peroxide. Int J Mol Sci 41:2847–2854.

30. Meinken C, Blencke HM, Ludwig H, Stülke J. 2003. Expression of the glycolytic *gapA* operon in *Bacillus subtilis:* differential syntheses of proteins encoded by the operon. Microbiology 149:751–761.

31. Nicolas P, Mäder U, Dervyn E, Rochat T, Leduc A, Pigeonneau N, Bidnenko E, Marchadier E, Hoebeke M, Aymerich S, Becher D, Bisicchia P, Botella E, Delumeau O, Doherty G, Denham EL, Fogg MJ, Fromion V, Goelzer A, Hansen A, Härtig E, Harwood CR, Homuth G, Jarmer H, Jules M, Klipp E, Chat LL, Lecointe F, Lewis P, Liebermeister W, March A, Mars RAT, Nannapaneni P, Noone D, Pohl S, Rinn B, Rügheimer F, Sappa PK, Samson F, Schaffer M, Schwikowski B, Steil L, Stülke J, Wiegert T, Devine KM, Wilkinson AJ, Dijl JM van, Hecker M, Völker U, Bessières P, Noirot P. 2012. Condition-dependent transcriptome reveals high-level regulatory architecture in *Bacillus subtilis*. Science 335:1103–1106.

32. Belitsky BR, Sonenshein AL. 1997. Altered transcription activation specificity of a mutant form of *Bacillus subtilis* GltR, a LysR family memner. J Bacteriol 179:1035–1043.

33. Martin-Verstraete I, Débarbouillé M, Klier A, Rapoport G. 1994. Interactions of wild-type and truncated LevR of *Bacillus subtilis* with the upstream activating sequence of the levanase operon. J Mol Biol 241:178–192.

34. Lachowicz TM, Morzejko E, Panek E, Piątkowski J. 1996. Inhibitory action of serine on growth of bacteria of the genus *Bacillus* on mineral synthetic media. Folia Microbiol 41:21–25.

35. Lamb DH, Bott KF. 1979. Inhibition of *Bacillus subtilis* growth and sporulation by threonine. J Bacteriol 137:213–220.

36. Wirtz L, Eder M, Brand AK, Jung H. 2021. HutT functions as the major L-histidine transporter in *Pseudomonas putida* KT2440. FEBS Lett 595:2113–2126.

37. Park JH, Lee KH, Kim TY, Lee SY. 2007. Metabolic engineering of *Escherichia coli for* the production of L-valine based on transcriptome and in silico knockout simulation. Proc Natl Acad Sci U S A 104:7797–7802.

38. Kennerknecht N, Sahm H, Yen MR, Patek M, Saier MH, Eggeling L. 2002. Export of L-isoleucine from *Corynebacterium glutamicum:* a two-gene-encoded member of a new translocator family. J Bacteriol 184:3947–3956.

39. Eggeling L, Sahm H. 2003. New ubiquitous translocators: amino acid export by *Corynebacterium glutamicum* and *Escherichia coli*. Arch Microbiol 180:155–160.

40. Brinkman AB, Ettema TJG, de Vos WM, van der Oost J. 2003. The Lrp family of transcriptional regulators. Mol Microbiol 48:287–294.

41. Schnetz K, Rak B. IS5: a mobile enhancer of transcription in *Escherichia coli*. Proc Natl Acad Sci U S A 89:1244–1248.

42. Sambrook J, Fritsch EF, Maniatis T. 1989. Molecular cloning: a laboratory manual, 2nd ed. Cold Spring Harbor Laboratory, Cold Spring Harbor, N.Y.

43. Kunst F, Rapoport G. 1995. Salt stress is an environmental signal affecting degradative enzyme synthesis in *Bacillus subtilis*. J Bacteriol 177:2403–2407.

44. Barbe V, Cruveiller S, Kunst F, Lenoble P, Meurice G, Sekowska A, Vallenet D, Wang T, Moszer I, Médigue C, Danchin A. 2009. From a consortium sequence to a unified sequence: the *Bacillus subtilis* 168 reference genome a decade later. Microbiology 155:1758–1775.

45. Kearse M, Moir R, Wilson A, Stones-Havas S, Cheung M, Sturrock S, Buxton S, Cooper A, Markowitz S, Duran C, Thierer T, Ashton B, Meintjes P, Drummond A. 2012. Geneious basic: an integrated and extendable desktop software platform for the organization and analysis of sequence data. Bioinformatics 28:1647–1649.

46. Reuß DR, Faßhauer P, Mroch PJ, Ul-Haq I, Koo BM, Pöhlein A, Gross CA, Daniel R, Brantl S, Stülke J. 2019. Topoisomerase IV can functionally replace all type 1A topoisomerases in *Bacillus subtilis*. Nucleic Acids Res 47:5231–5242.

47. Diethmaier C, Newman JA, Kovács AT, Kaever V, Herzberg C, Rodrigues C, Boonstra M, Kuipers OP, Lewis RJ, Stülke J. 2014. The YmdB phosphodiesterase is a global regulator of late adaptive responses in *Bacillus subtilis*. J Bacteriol 196:265–275.

48. Weinrauch Y, Msadek T, Kunst F, Dubnau D. 1991. Sequence and properties of *comQ*, a new competence regulatory gene of *Bacillus subtilis*. J Bacteriol 173:5685–5693.

49. Mehne FMP, Gunka K, Eilers H, Herzberg C, Kaever V, Stülke J. 2013. Cyclic di-AMP homeostasis in *Bacillus subtilis:* both lack and high level accumulation of the nucleotide are detrimental for cell growth. J Biol Chem 288:2004–2017.

50. Schilling O, Frick O, Herzberg C, Ehrenreich A, Heinzle E, Wittmann C, Stülke J. 2007. Transcriptional and metabolic responses of *Bacillus subtilis* to the availability of organic acids: transcription regulation is important but not sufficient to account for metabolic adaptation. Appl Environ Microbiol 73:499–507.

51. Ludwig H, Meinken C, Matin A, Stülke J. 2002. Insufficient expression of the *ilv-leu* operon encoding enzymes of branched-chain amino acid biosynthesis limits growth of a *Bacillus subtilis ccpA* mutant. J Bacteriol 184:5174–5178.

52. Kohlstedt M, Sappa PK, Meyer H, Maaß S, Zaprasis A, Hoffmann T, Becker J, Steil L, Hecker M, van Dijl JM, Lalk M, Mäder U, Stülke J, Bremer E, Völker U, Wittmann C. 2014. Adaptation of *Bacillus subtilis* carbon core metabolism to simultaneous nutrient limitation and osmotic challenge: a multi-omics perspective. Environ Microbiol 16:1898–1917.

53. Guder JC, Schramm T, Sander T, Link H. 2017. Time-optimized isotope ratio LC-MS/MS for high-throughput quantification of primary metabolites. Anal Chem 89:1624–1631.

54. Ludwig H, Homuth G, Schmalisch M, Dyka FM, Hecker M, Stülke J. 2001. Transcription of glycolytic genes and operons in *Bacillus subtilis:* evidence for the presence of multiple levels of control of the *gapA* operon. Mol Microbiol 41:409–422.

